# Rac1 contributes to brain connectivity impairments and neuropsychiatric disorders in Tuberous Sclerosis Complex

**DOI:** 10.64898/2025.12.09.693152

**Authors:** Tomasz Dulski, Olga Doszyń, Justyna Zmorzyńska

## Abstract

Tuberous sclerosis complex (TSC) is a neurodevelopmental disorder caused by inactivating mutations in the *TSC1* or *TSC2* genes, leading to constitutive activation of the mTOR complex 1 pathway. Patients commonly present with epilepsy and TSC-associated neuropsychiatric disorders (TANDs), including intellectual disability and anxiety. Our previous work identified anxiety-like behavior in *tsc2^vu242^* zebrafish mutants, characterized by enhanced thigmotaxis. We found that anxiety-like behavior in *tsc2^vu242^* zebrafish mutants was rescued by Rac1 inhibitors. These findings prompted further investigation into Rac1 signaling in the context of TSC pathology. Using G-LISA and FRET live imaging, we confirmed elevated Rac1 activity in the mutant brains. Rac1 activity was reduced following NSC23766 and EHT1864 treatment. Compared to wild-type controls, Rac1 activity in mutants was elevated by 50% in untreated animals, and reduced by ∼30–40% after Rac1 inhibitors. Moreover, aberrant connectivity between brain hemispheres that correlated with anxiety before was restored after Rac1 inhibition. Transcriptomic profiling revealed 1,260 DEGs between homozygous and heterozygous *tsc2^vu242^* zebrafish. Rac1 inhibition restored the expression of approximately 1,000 genes toward wild-type levels. KEGG pathway analysis identified significant enrichment in mTOR signaling and focal adhesion, while GO terms highlighted processes including axon guidance, axonogenesis, and neuronal projection. Collectively, our findings support a functional role for Rac1 in TSC pathology, linking cytoskeletal dysregulation with aberrant axon development and anxiety-like behavior. Rac1 inhibition emerges as a promising therapeutic avenue for modulating both brain structure and function in TSC-associated anxiety.

**Highlights:**

- Rac1 activity is upregulated in *tsc2^vu24/vu242^* zebrafish mutants and correlates with anxiety-like behavior.
- Inhibition with NSC23766 and EHT1864 reduces Rac1 activity, partially normalizes mTORC1 signaling, and rescues anxiety-like behavior.
- Live FRET imaging and G-LISA confirm elevated Rac1 signaling and its modulation by pharmacological treatment.
- Transcriptomic profiling reveals 1,260 deferentially expressed genes in mutants, with restoration of ∼1,000 genes by NSC23766.
- Gene expression and pathway analysis link Rac1 dysregulation to altered actin dynamics, axon guidance, and focal adhesion.

## 1. Introduction

Zebrafish have emerged as a valuable *in vivo* model for studying early neurodevelopmental processes due to their external development, optical transparency, and genetic tractability. The *tsc2^vu242^* zebrafish mutant line recapitulates key features of Tuberous sclerosis complex (TSC), including white matter disorganization and behavioral abnormalities (Kedra et al., 2020; Kim et al., 2011). TSC is an autosomal disease caused by inactivating mutations in either the *TSC1* or *TSC2* gene. The protein products of these genes, Hamartin and Tuberin, form a complex that negatively regulates the activity of mechanistic/mammalian target of rapamycin complex 1 (mTORC1) (Huang et al., 2008). mTORC1 is a key integrator of growth and signaling cues, and plays critical roles in neuronal development, including axon and dendrite formation, synaptic plasticity, and network connectivity (Klan et al., 2004). In the brain, TSC manifests as cortical dysplasia, epilepsy, and a spectrum of neuropsychiatric symptoms collectively referred to as TSC-associated neuropsychiatric disorders (TANDs), which include intellectual disability, autism spectrum disorder, attention-deficit/hyperactivity disorder, and anxiety (de Vries et al., 2018). TANDs do not fully correlate with mTOR expression levels or epilepsy, suggesting that other molecular pathways may interact with mTORC1 to produce these symptoms.

Previously, we observed elevated *rac1*, *dock4* and *elmo2* transcript levels in *tsc2^vu242/vu242^* larvae, suggesting hyperactivation of Rac1 (Kedra et al., 2020). Rac1 is a small GTPase that regulates actin remodeling and neuronal morphology by activation through interactions with guanine nucleotide exchange factors (GEFs) such as ELMO and DOCK proteins, which facilitate the transition to its active, GTP-bound form (Kedra et al., 2020). In this paper, we link *tsc2* loss, Rac1 overactivation, with dysregulation of actin-related genes and altered brain connectivity, which leads to anxiety-like behavior. Our data support the role of Rac1 signaling as in the pathogenesis of TSC and suggest its inhibition as a promising therapeutic strategy for alleviating neurodevelopmental symptoms associated with the disorder.

## 2. Materials and methods

### 2.1. Zebrafish

Heterozygous zebrafish (*Danio rerio*) carrying the *tsc2^vu242/+^* mutation were used. The offspring were raised to adulthood and genotyped by high resolution melting (HRM) technique. Subsequently, *tsc2^vu242/+^* heterozygous were selected and in-crossed. The *tsc2^+/+^, tsc2^vu242/+^* and *tsc2^vu242/vu242^* F2 generation larvae were produced in a Mendelian ratio of 1:2:1, identified by HRM, and used for the experiments.

### 2.2. Zebrafish husbandry

Zebrafish adults were maintained at 28,5 ± 0,5°C on a 14:10 h light/dark cycle under standard aquaculture conditions. Embryos were collected via natural spawning and immediately transferred to embryo medium (1.5 mM HEPES buffer (pH 7.2), 17.4 mM NaCl, 0.21 mM KCl, 0.18 mM Ca(NO3)2, 0.12 mM MgSO4) and 0.6 µM methylene blue. For all experiments, zebrafish larvae aged 0–10 days post-fertilization (dpf) were used and maintained in an incubator under a 14 h light / 10 h dark photoperiod at 28.5 °C, in accordance with European guidelines for the protection of animals used for scientific purposes (Directive 2010/63/EU). Experiments involving animals beyond 120 hpf were conducted under approval from the institutional animal ethics committee, as required for procedures performed on free-feeding zebrafish larvae.

### 2.3. Drug treatments

Zebrafish embryos after fertilization were collected and placed into a petri dish (50 per dish), containing 40 ml of E3 embryo medium. Tested compounds (NSC23766, Tocris, cat. No. 2161; EHT1864, Tocris, cat. No: 3872) were diluted in E3 medium. Rapamycin (Medchem Express, cat. No. HY-10219) superstock (1mM) was first diluted in DMSO and further dilutions were done in E3. In most experiments 1 uM NSC23766 and 50 nM EHT1864 were used and added directly to E3 medium at 96 hpf. In case of rapamycin, 200 nM concentration was used starting at 48 hpf with daily exhcange in order to prevent disease-asosciated phenotypes appearance (RapaP).

### 2.4. Genotyping

For genotyping zebrafish were euthanized in a solution of tricaine methanesulfonate (MS222, 0.765 mmol/L). Depending on the experiment performed, a fin clip, the trunks or whole larvae were used for the extraction of the genomic DNA. The samples were digested in 20 µl E3 with 3 mg/ml of proteinase K by heating them for 3h at 55°C. Afterwards, the proteinase K was inactivated for 10 min at 95°C. For genotyping, 1 μl of genomic DNA was mixed with 5 μL of the RT PCR Mix EvaGreen for HRM analysis (A&A Biotechnology cat: 2008-1000G), 0.5 μl of each primer (10 μM) and 3 µl milliQ water. The following primers were used; GAGACC TGCCTGGACATGAT (*tsc2*, forward primer) and CTTGGGCAGAGCAGAGAAGT (*tsc2*, reverse primer). The HRM reaction was performed in the Lightcycler 96 instrument (Roche, cat: 05815916001) using dedicated Software for this machine (Roche).

### 2.5. Vector cloning

The mCerulean-Rac1-Ypet (Rac1-Raichu) fusion construct was generated using the Sequence and Ligation Independent Cloning method. The Rac1 coding region and associated regulatory domains were amplified from the pEYFP-Rac1-ECFP plasmid (known as Raichu-Rac1/Rac1-CT) (Yoshizaki et al., 2003), kind gift from Prof. Michiyuki Matsuda, Kyoto University). Next fluorescent protein genes mCerulean and Ypet were amplified from the pcDNA3-TorCar (#64927 Addgene, kind gift from Prof. Jacek Jaworski, IIMCB) plasmid and inserted upstream and downstream of Rac1, respectively. The resulting mCerulean-Rac1-Ypet sequence was cloned into the pCS2+ expression vector (kind gift from Professor William A Harris, Cambridge University) for downstream applications. All cloning steps were verified by Sanger sequencing.

### 2.6. Electroporation of plasmid into the zebrafish brains & FRET in vivo for Rac1 activity measurement

Zebrafish embryo were microinjected with mCerulean-Rac1-Ypet plasmid. Due to high mortality of the embryos after microinjection to the cell at one-cell stage of fertilized zebrafish egg, we microinjected the plasmid at 48 hpf to brain ventricle and electroporated it into the right brain hemisphere. For microinjections, the micropipettes were heat-pulled from borosilicate capillaries (I.D 0.75mm O.D. 1.0 mm) and loaded with the plasmid DNA solution in 0.1M KCl containing 0.05% Phenol Red and 25ng of DNA. At 24hpf, fish were immobilized in a drop of low melting point agarose. We mounted the fish as previously reported (Doszyn et al., 2025). After the injections, the fish were immediately electroporated for plasmid migration into the brain cells (NEPA21 electroporator, Nepa gene, Japan). Then fish were raised in the normal light cycle. The embryos were imaged *in vivo* using single-plane illumination microscopy (SPIM) with a Lightsheet Z.1 microscope, and afterwards clipped for genotyping. Fluorescence resonance energy transfer (FRET) was used to detect Rac1 conformational change upon activation by GEFs. FRET efficiency was assessed using a donor-dequenching approach. Cells expressing the mCerulean-Rac1-Ypet plasmid were first imaged under CFP excitation and YFP emission spectrum acquisition to capture the initial FRET signal, followed by imaging of CFP and YFP spectra separately. YFP was then selectively photobleached to eliminate acceptor fluorescence. A second set of images was subsequently collected under the same conditions, allowing detection of the increase in donor emission resulting from loss of FRET. FRET efficiency was calculated from the relative change in donor signal before and after photobleaching and corrected for bleaching efficiency.

### 2.7. G-LISA Rac1 activity measurement

For Rac1 activity measurement, G-LISA was performed according to the manufacturer’s protocol (cat. no. BK128, Cytoskeleton Inc.). Rac1 Activation Assay is a commercially available enzyme linked immunosorbent assay (ELISA)-based method which allows measuring GTP-bound (active) Rac1 in lysed samples. It utilizes a 96-well plate coated with Rac1-binding domain of PAK, the Rac1 effector, that binds only GTP-bound form (active) but not the inactive GDP-bound form of Rac1. Detection was made by specific primary and HRP-conjugated secondary antibodies provided in the kit.

### 2.8. Image acquisition & analyses

Images of fluorescently labeled whole mounts immunostainings and in vivo imaging of neuronal activity in the zebrafish brains were acquired using a Lighsheet Z.1 microscope (40× objective, NA = 1.3) and an incubation chamber with temperature control. For the imaging, Z-stacks were taken with an optimal interval of 0.48 μm. Analyses were performed using Fiji software (Schindelin et al., 2012). Regions of interest (ROIs) were drawn manually. Cells in WM were calculated automatically on thresholded images using the Particle Analysis tool. Cell size, fluorescence intensity, and commissural width and volume were measured using the measurement tool.

### 2.9. Behavioral analyses

Before the measurements, 5 dpf larvae were acclimated into the behavioral room for 5 min. The basal locomotor activity test was performed in 24-well plates after overnight habituation of the larvae. Experiments were performed in a total volume of 0.5 ml. The recordings were taken in the dark for 1 h, with a 600 s data integration period for basal locomotor activity, and for 30 min, with a 5 s data integration period and a final light power of 30% during the light phase (bottom source of light) for the dark phase. The experimental setup included equal 10 min light and then dark periods. Open field test is already well established in our lab and was performed as previously described (Kedra et al., 2020).

### 2.10. Immunoblotting

Protein extraction and immunoblotting were performed according to a standard protocol (zfin.org). Briefly, a fresh, cold Ringer buffer (supplemented with ethylenediaminetetraacetic acid and phosphatase and protease inhibitors) was applied for washing and storing. Sodium dodecyl sulfate (SDS) sample buffer was used for protein extraction from at least 20 heads per genotype pulled in the sample. Lysates were loaded on polyacrylamide gels, run by SDS-polyacrylamide gel electrophoresis, and transferred to nitrocellulose membranes. The following antibodies were used: Rac1/2/3 (1:400, cat. No. sc-514583, Santa Cruz), anti-Tubulin (1:2000, catalog no. T516B, Sigma-Aldrich), Peroxidase AffiniPure Donkey Anti-Rabbit IgG (H+L) (1:10 000, cat. No. 711-035-152, Jackson Immunoresearch), Peroxidase AffiniPure Donkey Anti-Mouse IgG (H+L) (1:10 000, cat. No. 715-035-150, Jackson Immunoresearch). Membranes were analyzed using the Amersham ImageQuant 800 (Cytiva).

### 2.11. Whole-Mount Immunostainings

Whole-mount antibody staining (WMAS) were performed according to the protocol described elsewhere (Doszyn et al., 2025) The following antibodies were used: anti-Tubulin (Acetyl LYS40), (1:400, cat. No. GTX16292, Genetex), anti-P-S235/236-Rps6 (1:400, catalog no. CS4858, Cell Signaling Technology).

### 2.12. RNA sequencing and sample preparation

*Tsc2 ^+/+^*, *tsc2 ^vu242/+^* and *tsc2 ^vu242/vu242^* zebrafish larvae (5dpf) were euthanized by immersion in MS222, 0.765 mmol/L. Fish were decapitated with a scalpel. The tails were used for genotyping. Heads were used for brain extraction. 40 extracted brains were pooled (per genotype and treatment) and RNA extractions were performed using Monarch® Total RNA Miniprep Kit (New England Biolabs, cat. T2010S) according to the manufacturer’s instructions. To purify samples, Monarch® RNA Cleanup Kit (New England Biolabs, cat. T2040L) was used. The concentration and purity of the purified total RNA were determined spectrophotometrically using the Nanodrop. High Sensitivity RNA analyses were done on the Agilent 2200 TapeStation instrument to assess RNA integrity. 500 ng of total RNA per sample with a RIN score above 7 was taken into consideration for library prep. Isolation of poly(A)+ RNA transcript from Total RNA for RNA library preparation was performed using NEBNext® Poly(A) mRNA Magnetic Isolation Module (NEB #E7490S/L). The NEBNext® Ultra II Directional RNA Library Prep Kit for Illumina® kit was used for library preparation. Sequencing was performed thanks to Genomics Core Facility CeNT UW (Europe, Poland). The library was subjected to paired-end reading mode 2×100 cycles with sequencing using the Novaseq6000 platform with the standard procedure recommended by the manufacturer.

### 2.13. Bioinformatic analysis

Read quality was checked using FastQC v 0.11.9 software (Andrews et al., 2010). Low-quality reads were filtered out using Trimmomatic v 0.38 (Bolger et al., 2014): low-quality leading and trailing bases were removed, and the quality of the body of the reads was assessed with a trimming sliding window of 4, and a Phred score threshold of 20 bases. Reads that passed quality control were aligned to the zebrafish reference genome, GRCz11, using Salmon v 1.4.0 (Patro et al., 2017). The number of reads that align to each gene in accordance with the Ensembl GRCz11 zebrafish reference annotation was around 80%. Differential expression analysis was carried out using the R package DESeq2 (Love et al., 2014). Gene expression changes with a Benjamini-Hochberg adjusted p-value < 0.05 were considered statistically significant. For comparing the transcriptomes of the zebrafish mutants heterozygous and wildtypes in different treatments, zebrafish identifiers of genes differentially expressed in our larvae comparisons were converted to their human orthologs using BiomaRt (Durinck et al., 2009).

### 2.14. Gene ontology and KEEG pathway enrichment analysis

The Kyoto Encyclopedia of Genes and Genome (KEGG) (Kanehisa et al., 2012) for zebrafish was queried to test differentially expressed genes (DEGs) in zebrafish larvae for pathway enrichment, respectively, using the enrichKEGG function from clusterProfiler package v 4.6.2 (Wu et al. 2021). The Gene Ontology (GO) knowledgebase (Ashburner et al., 2000) for zebrafish was queried to test DEGs in zebrafish larvae for gene ontology enrichment, respectively, using the enrichGO function from clusterProfiler package. Enriched GO term lists containing the top terms amongst biological processes were processed to create a visual representation of the GO enrichment.

### 2.15. Statistical analyses and experimental design

No predetermination of sample sizes was performed because the number of each genotype could not be predicted due to the random distribution and early lethality of homozygotes. The data were plotted to verify normality. The normality of data distributions was assessed using the Shapiro-Wilk test applied to model residuals. Homogeneity of variance was evaluated using Levene’s test. When assumptions of normality and equal variance were met, data were analyzed using one-way or two-way analysis of variance (ANOVA) (depending on the number of factors) and followed by Tukey’s Honestly Significant Difference (TukeyHSD) for multiple-comparison adjustments. When these assumptions were violated or when sample sizes were too small to reliably assess normality, nonparametric tests were applied. Specifically, the Kruskal–Wallis test was used for multi-group comparisons, and pairwise group differences were further examined using the Wilcoxon rank-sum test. For Wilcoxon tests, p-values were adjusted using the Benjamini–Hochberg false discovery rate (FDR) method. An adjusted p < 0.05 was considered statistically significant. Statistical significance in figures was denoted as follows: *p < 0.05, **p < 0.01, ***p < 0.005, ****p < 0.001. All the analyses were performed using RStudio. Data are presented as medians with Q1 and Q3 quartiles using boxplots, unless otherwise stated. Open dots represent data points. Filled dots represent data outliers. In most cases, randomization was assured by a lack of knowledge about the genotypes while performing the experiments and the automatic or semi-automatic analyses.

## 3. Results

### 3.1. Rac1 inhibitors NSC23766 and EHT1864 rescue anxiety-like behavior in tsc2^vu242/vu242^ zebrafish

We have previously shown that the *tsc2^vu242/vu242^* fish exhibit hypervelocity, thigmotaxis in the open field test, and anxiety-like behavior in response to a sudden change in light conditions (Kedra et al., 2020). Here, we have performed the open field using *tsc2^vu242^* fish pretreated with NSC23766 and EHT1864 versus untreated, and have confirmed a rescue of anxiety-like behaviors in those tests by both Rac1 inhibitors.

Before starting the experiments with treatments, we did the toxicity screening of Rac1 inhibitors to choose appropriate doses (Fig. 1A). For survival toxicity screening embryos were exposed continuously to the compounds from 24 hpf up to 240 hpf. Mortality and morphological abnormalities were assessed at 24, 48, 72, 96, 120, 144, 168, 192, 216 and 240 hpf under a stereomicroscope. Observed endpoints included delayed development, pericardial edema, tail malformation, yolk sac edema, and lack of spontaneous movement. Based on this screening, the selected working concentrations —1 µM NSC23766 and 50 nM EHT1864 — did not induce mortality, developmental delay, or morphological abnormalities, indicating that both doses were non-toxic for zebrafish across the full exposure window.

**Figure 1.**
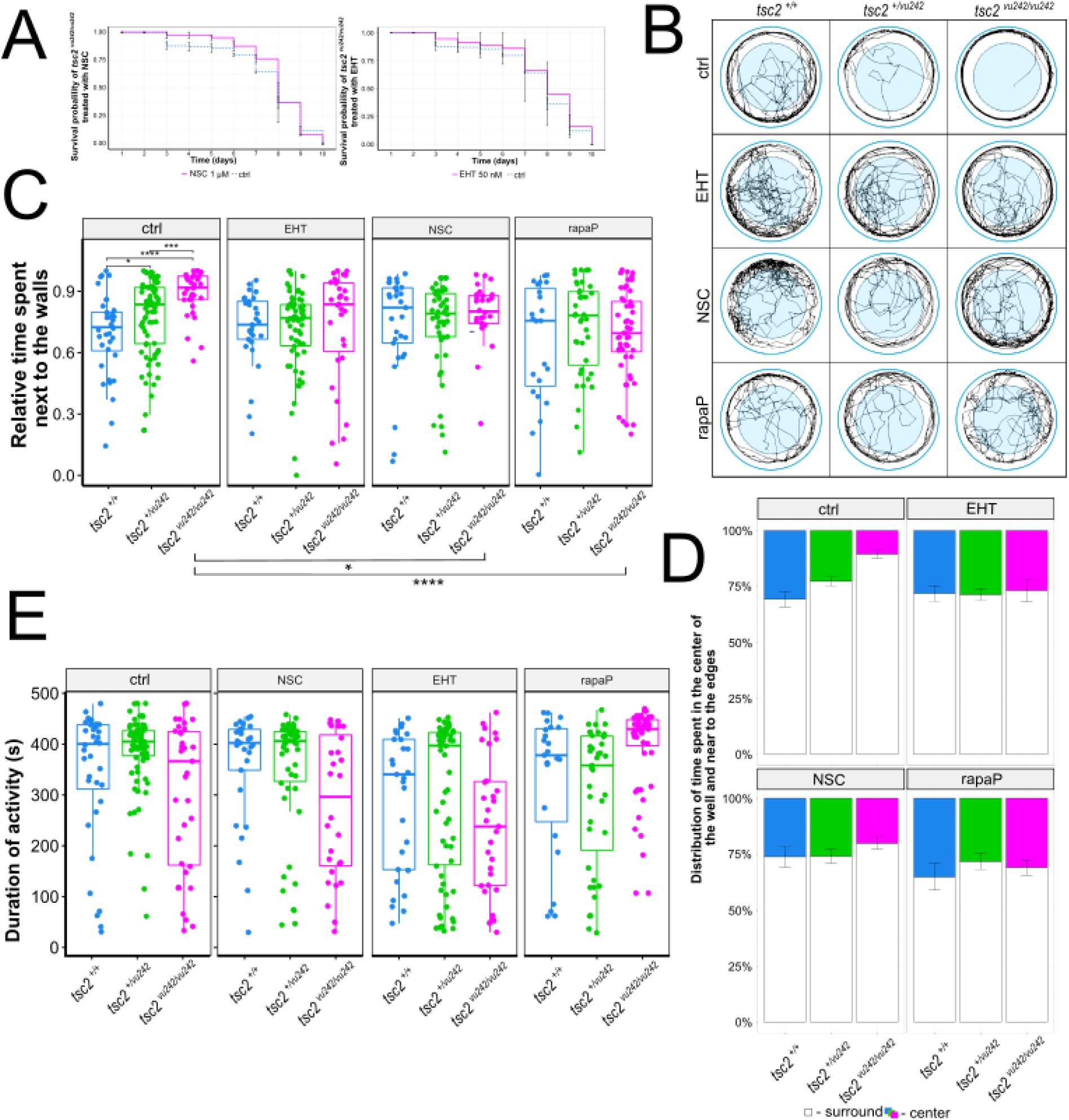
*Tsc2^vu242/vu242^* fish show anxiety-like behavior that is ameliorated by pretreatment with NSC23766 and EHT1864. RapaP – rapamycin pretreatment starting at 2 dpf in order to prevent the development of symptoms. (**A**) Toxicity screening of *tsc2^vu242/vu242^* treated with Rac1 inhibitors vs. control *tsc2^vu242/vu242^*. Chosen doses (1 μM NSC23766, 50 nM EHT1864) did not influence on *tsc2^vu242/vu242^* mortality. (**B**) Representative tracks from the open field test for each *tsc2^vu242^* genotype in the control group vs. treated with NSC23766 and EHT1864. Summary tracks from 8 min recordings show thigmotaxis behavior in *tsc2^vu242/vu242^* fish that is rescued by NSC23766 treatment. (**C**) Relative time spent next to the walls of the fish [p = 0.002 for control *tsc2^+/+^* vs. control *tsc2^vu242/vu242^*, p = 0.000159 for control *tsc2^vu242/+^* vs. control *tsc2^vu242/vu242^*, p = 0.001 for control *tsc2^vu242/vu242^* vs. NSC23766 *tsc2^vu242/vu242^*, p = 1.25 ×10^−6^ for control *tsc2^vu242/vu242^* vs. rapaP *tsc2^vu242/vu242^*]. (**D**) Distribution of time spent in the center and next to the walls of the dish of all fish across treatment groups. (**E**) Duration of overall activity across all genotype and treatment groups. No significant difference between median activity of each group indicates that the relative activity differences are not caused by changes in overall motor activity.

As expected, *tsc2^vu242/vu242^* larvae spent significantly more time near the walls of the testing area compared with wild-type (*tsc2^+/+^*) and heterozygous (*tsc2^vu242/+^*) siblings, consistent with increased thigmotaxis and anxiety-like behavior. This was reduced in *tsc2^vu242/vu242^*homozygotes following treatment with NSC23766 (Fig.1 B-E). In case of EHT1864 treatment, thigmotaxis was reduced partially for some fish. In parallel, a reduction in thigmotaxis was also observed following pretreatment with the RapaP (mTORC1 inhibitor rapamycin) (Fig.1 B-E).

### 3.2. NSC23766 and EHT1864 treatments affects Rac1 activation in tsc2^vu242/vu242^ zebrafish brains

To determine whether NSC23766 and EHT1864 modulate Rac1 activity in our model, we assessed Rac1 activation using G-LISA assays. Previous qPCR analyses revealed increased expression of *rac1*, along with elevated levels of *elmo2* and *dock3*, *dock4* transcripts in *tsc2^vu242/vu242^*larvae, suggesting upregulation of Rac1 pathway at the transcriptional level in *tsc2^vu242/vu242^* mutants (Kedra et al., 2020). G-LISA assays confirmed increased GTP-bound Rac1 levels detected in non-treated *tsc2^vu242/vu242^*vs. control *tsc2^+/+^*. Treatment with the both Rac1 inhibitors NSC23766 and EHT1864 reduced Rac1 activity in mutant larvae. NSC23766 had a stronger effect and restored Rac1 activity to near wild-type levels (Fig. 2D). Based on the Rac1 activity ratio relative to untreated *tsc2^vu242/vu242^* control, NSC23766 treatment reduced Rac1 activity by an average of ∼36% in mutants, while EHT1864 treatment resulted in a ∼25% reduction (Fig. 2D). In line with this Rac1 activity ratio to untreated *tsc2 ^+/+^*showed increased Rac1 activity in *tsc2^vu242/vu242^* control up to average of ∼46%, whereas in case of treated *tsc2* mutants with NSC23766 and EHT1864 Rac1 activity lower average of ∼8% and raised only up to ∼6 % respectively. Furthermore, to directly visualize Rac1 activation *in vivo*, we employed a FRET-based Rac1 biosensor (Raichu-Rac1 construct) (Fig. 2A) that was microinjected and electroporated (Fig.2 B-C). Increased FRET efficiency was observed in the brains of *tsc2^vu242/vu242^* compared to *tsc2^+/+^* siblings, indicating elevated Rac1 activity (Fig. 3 C). Immunoblotting showed no statistically significant differences in total Rac1 protein levels across all genotypes (Fig. 2E), indicating that the observed changes in activity stem from altered upstream regulation and active Rac1 level rather than expression.

**Figure 2.**
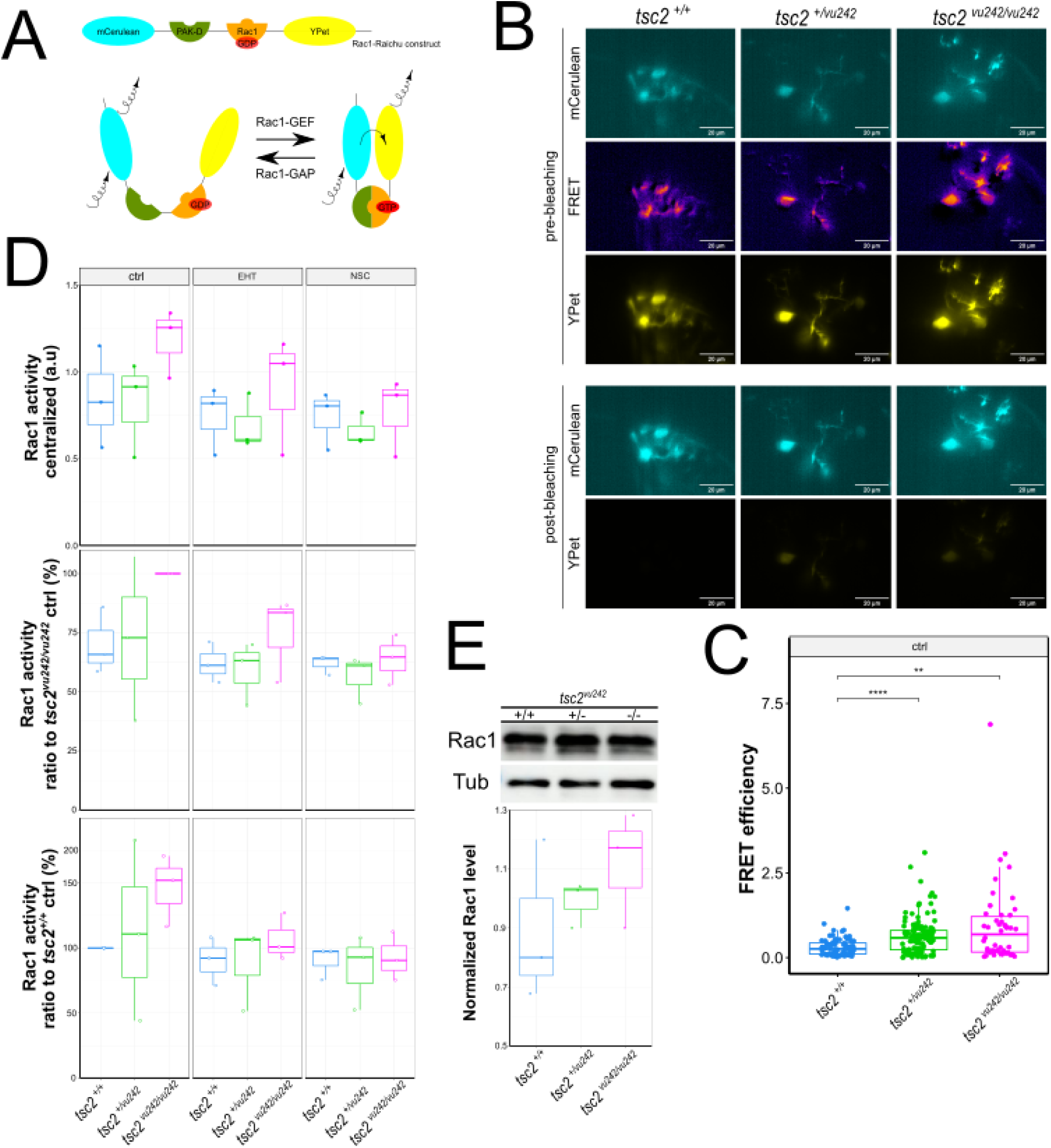
Characterization of Rac1 pathway activity in *tsc2^vu242^* zebrafish brain. (**A**) Structure of prepared Rac1-Raichu construct used in *tsc2^vu242/vu242^* zebrafish to test Rac1 activity in zebrafish brain by FRET efficiency. Figure presents the principle of the FRET technique. In brief, active Rac1 (GTP-bound) undergoes a conformational change that promotes binding to the PAK domain, bringing mCerulean (donor) and YPet (acceptor) into close proximity. This spatial arrangement enables energy transfer from mCerulean to YPet upon donor excitation. The resulting increase in YPet emission serves as a readout of Rac1 activation, allowing dynamic visualization of Rac1 activity in live cells. (**B**) FRET-based Rac1 activity biosensor imaging in control *tsc2^+/+^* vs. *tsc2^vu242/vu242^* zebrafish brains injected with mCerulean-Rac1-Ypet plasmid. (**C**) Quantification of FRET efficiency [padj = 0.002 for control *tsc2^+/+^* vs. control *tsc2^vu242/vu242^*, padj = 1.83 ×10^−7^ for control *tsc2^+/+^*vs. control *tsc2^vu242/+^*] (**D**) G-LISA quantification of Rac1 activity of all fish across genotypes and treatment groups. (**E**) Representative immunoblots and quantification of Rac1 protein levels for control *tsc2^vu242^* zebrafish.

**Figure 3.**
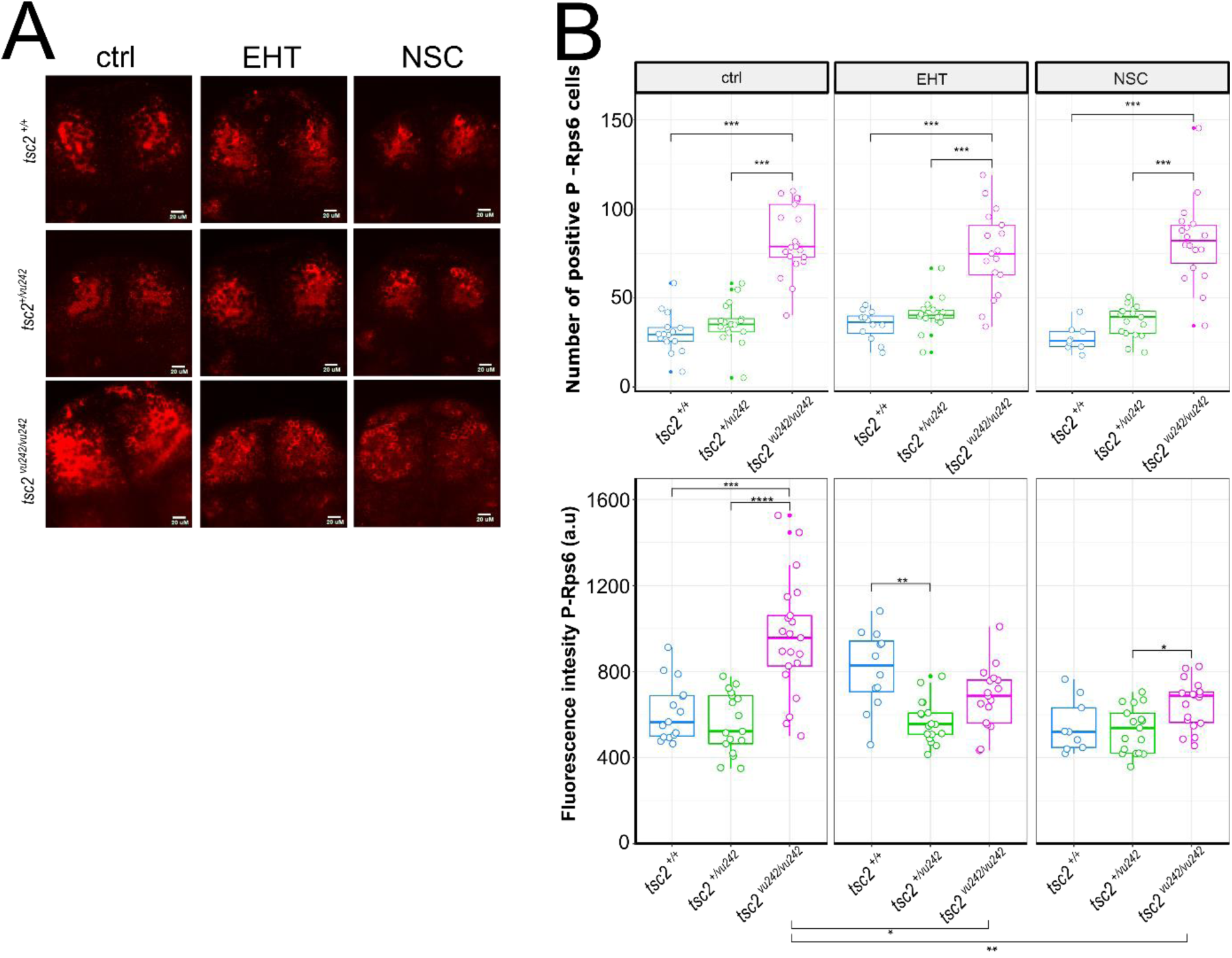
EHT1864 and NSC23766 modulates mTORC1 signaling in *Tsc2^vu242/vu242^* zebrafish. (A) WMAS images of the pallium stained with phosphorylated Rps6 levels in all genotypes of *tsc2^vu242^*fish with or without inhibitors treatment. Scale bar = 20 μm. (B) Quantification of numbers of positive P-Rps6 cells and mean P-Rps6 signal intensity in the pallium across genotypes ± EHT1864 and NSC23766. Each dot represents one individual fish. [No. of positive P-Rps6 cells - padj = 0.0000714 for control *tsc2^+/+^* vs. control *tsc2^vu242/vu242^*, padj = 0.00000981 for control *tsc2^vu242/+^*vs. control *tsc2^vu242/vu242^*, padj = 0.00192 for EHT1864 *tsc2^+/+^*vs. EHT1864 *tsc2^vu242/vu242^*, padj = 0.03 for EHT1864 *tsc2^vu242/+^* vs. EHT1864 *tsc2^vu242/vu242^,* padj = 0.00192 for NSC23766 *tsc2^+/+^* vs. NSC23766 *tsc2^vu242/vu242^*, padj = 0.00101 for NSC23766 *tsc2^vu242/+^*vs. NSC23766 *tsc2^vu242/vu242^*, Fluorescence Intensity - padj = 0.000748 for control *tsc2^+/+^* vs. control *tsc2^vu242/vu242^*, padj = 0. 0.0000354 for control *tsc2^vu242/+^* vs. control *tsc2^vu242/vu242^*, padj = 0.00650 for EHT1864 *tsc2^+/+^* vs. EHT1864 *tsc2^vu242/+^*, padj = 0.033 for NSC23766 *tsc2^vu242/+^* vs. NSC23766 *tsc2^vu242/vu242^*, padj = 0.010341 for EHT1864 *tsc2^vu242/vu242^* vs. control *tsc2^vu242/vu242^*, padj = 0.0015283 for NSC23766 *tsc2^vu242/vu242^*vs. control *tsc2^vu242/vu242^*].

**Figure 4.**
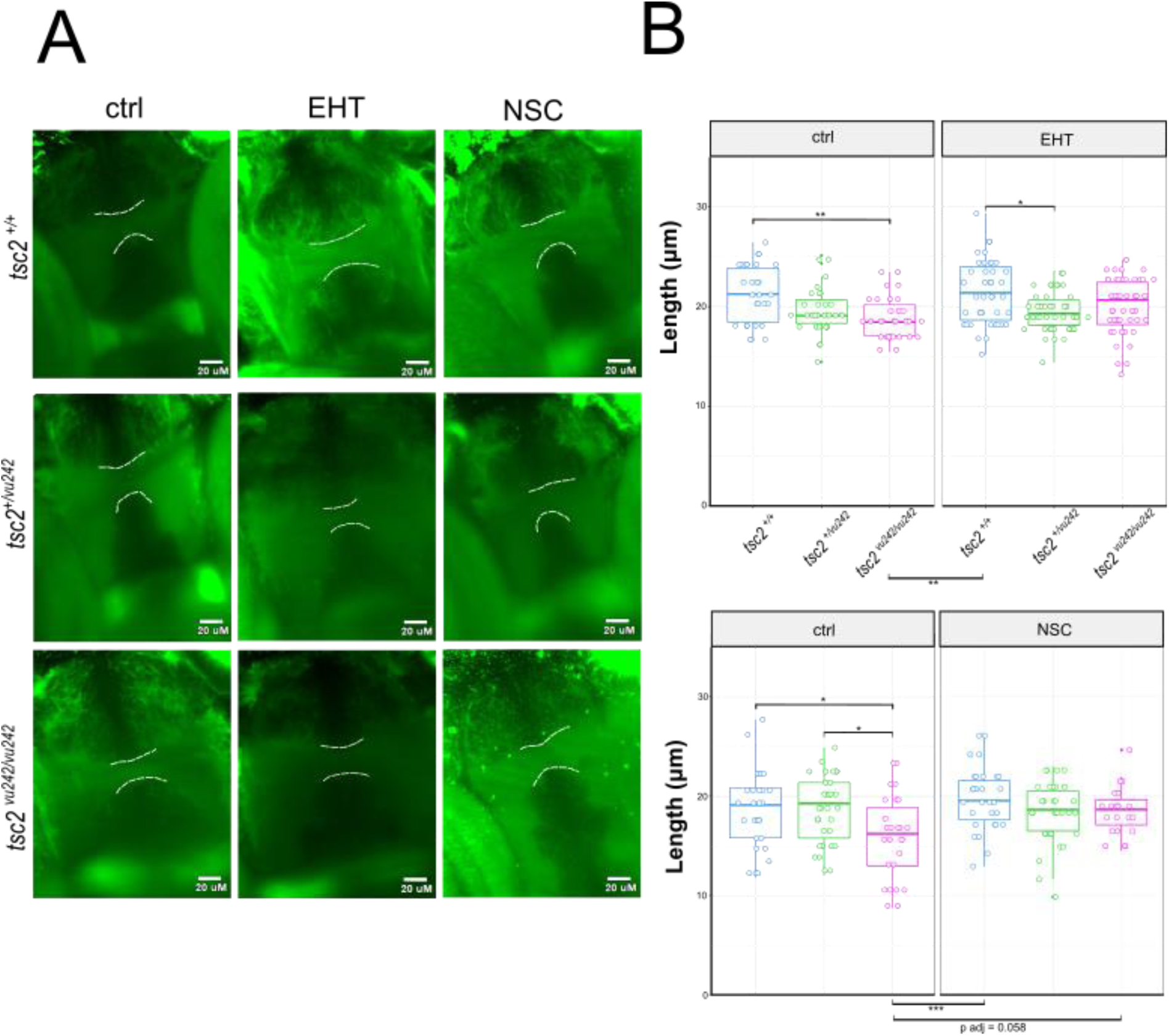
Rac1 pathway regulates axonal specification and connectivity in Tsc2-deficient zebrafish. (**A**) Immunostaining of AC axons tract formation under Rac1 inhibitors. WMAS images of the AC axons tract formation stained with Acetylated Tubulin (AcTub) in all genotypes of *tsc2^vu242^* fish with or without inhibitors treatment. Scale bar = 20 μm. (**B**) Quantification of AC axons width across genotypes ± EHT1864 and NSC23766. Each dot represents one individual fish. [Experiment with EHT1864 treatment - padj = 0.0352 for control *tsc2^+/+^* vs. control *tsc2^vu242/vu242^*, padj = 0.0930 for EHT1864 *tsc2^+/+^* vs. EHT1864 *tsc2^vu242/+^*, padj = 0.0084 for EHT1864 *tsc2^+/+^* vs. control *tsc2^vu242/vu242^*; Experiment with NSC23766 treatment-padj = 0.0338955 for control *tsc2^+/+^*vs. control *tsc2^vu242/vu242^,* padj = 0.0265729 for control *tsc2^vu242/+^* vs. control *tsc2^vu242/vu242^,* padj = 0.0005594 for NSC23766 *tsc2^+/+^* vs. control *tsc2^vu242/vu242^*]

### 3.3. Rac1 inhibition affects mTORC1 activity

Rac1 is a key regulator of cytoskeletal dynamics, but emerging evidence also links its activation to the modulation of mTORC1 signaling through PI3K-dependent and PI3K-independent pathways (Saci et al., 2011). Given that *tsc2^vu242/vu242^*zebrafish exhibit hyperactivation of mTorC1 due to loss of Tsc1-Tsc2 complex function, we tested whether inhibition of Rac1 by NSC23766 or EHT1864 could impact mTorC1 signaling as part of their therapeutic action. To evaluate mTorC1 activity, we performed WMAS against phosphorylated ribosomal protein S6 (P-Rps6), a well-established downstream target of mTorC1. As expected, untreated *tsc2^vu242/vu242^*mutants showed a significant increase in P-Rps6 compared to wild-type and heterozygous siblings, consistent with constitutive mTorC1 activation (Fig. 3 A-B). While high levels and broad distribution of P-Rps6-positive cells were observed in the pallium of untreated *tsc2^vu242/vu242^*, NSC23766 and EHT1864 treatments resulted in statistically significant reduction of the fluorescence intensity within these cells, suggesting decreased mTorC1 activity per cell. However, the total number of P-Rps6-positive cells did not significantly decrease following treatment (Fig. 3 A–B), indicating that Rac1 inhibition affects the magnitude of mTorC1 signaling rather than the extent of its activation across the tissue.

### 3.4. Effects of Rac1 Pathway Modulation on Neural Connectivity in Tsc2-Deficient Zebrafish

The anterior commissure (AC), a major forebrain commissural tract that connects the brain hemispheres in the telencephalon, is involved in interhemispheric communication and is sensitive to disruptions in neurodevelopmental signaling pathways (Fenlon et al., 2021). Our previous study showed that AC in our *tsc2^vu242/vu242^* is thinner than *in tsc2^+/+^* fish consistent with altered axon growth in the mutant brain (Kedra et al., 2020). Treatment with Rac1 inhibitors produced a modest increase in AC thickness in *tsc2^vu242/vu242^*, trending toward wild-type dimensions. NSC23766 treatment had a stronger effect on increasing AC thickness than EHT1864. Although this rescue was not fully statistically significant (padj = 0.056 for ^tsc2vu242/vu242^ treated with NSC23766 *vs*. untreated), it suggests that abnormal Rac1 signaling contributes to structural forebrain abnormalities in *tsc2^vu242/vu242^* fish.

### 3.5. Crosstalk Between Rac1 and Other Pathways in TSC Pathogenesis

Next, we performed transcriptomic analysis and integrative pathway mapping in the *tsc2^vu242/vu242^* zebrafish compared to wild-type siblings with and without treatment with NSC23766. Each sample consists of 40 brains extracted and pulled together. At least three replicates were collected for each genotype from each drug treatment and control. RNA-seq analysis of *tsc2^vu242/vu242^* zebrafish brains compared to wild-type (*tsc2^+/+^*) identified 1,260 DEGs from which 524 genes were downregulated and 736 genes were upregulated (Fig. 5A). Differential expression analysis between *tsc2^vu242/vu242^* and *tsc2^+/+^* identified subsets of zebrafish transcripts which were mapped to their corresponding human orthologs to enable cross-species pathway interpretation. For human orthologous DEGs, we found associations with Axon Guidance, Actin Cytoskeleton Dynamics, mTOR signaling, and Rac1 pathway regulation were the most significant. GO enrichment analyses of the upregulated DEGs between *tsc2 ^vu242/vu242^*zebrafish brains and *tsc2^+/+^* revealed significant enrichment in 17 genes related to neurodevelopmental pathways such as axon development or guidance, neuron projection morphogenesis or guidance (Fig. 5B). Furthermore, KEGG pathway analysis of the same set of genes revealed significant enrichment in actin cytoskeleton regulation, focal adhesion, mTOR related pathways (Lysosome, Autophagy, Motor proteins, Phagosomes, and Apoptosis that can all be related to mTOR activity, though in different ways). KEGG enrichment analysis also revealed mTor signaling pathway which confirmed involvement of mTor in our Tsc2-loss zebrafish model (Fig. 5 C). Detailed pathview of mTOR in *tsc2 ^vu242/vu242^*shows many dysregulations of upstream targets of the mTOR (Fig. 5F). We found overexpression of *tsc1b*, an isoform of Tsc1 protein which forms a functional complex with Tsc2 that negatively regulates mTorC1 activity. This overexpression might be associated with compensatory mechanisms of the cells. The network diagram illustrated that most of the DEGs of control *tsc2^vu242/vu242^*cluster together for mTOR related pathways (Lysosome, Autophagy, Motor proteins, Phagosomes, and Apoptosis, mTOR pathway) and Focal adhesion, ECM-receptor interaction. However, there are no shared molecular interactors linking mTOR and actin cytoskeleton signaling (Fig. 5 D). The overlap analysis identified 261 DEGs shared between the control Tsc2-deficient zebrafish brains and zebrafish brains (mutants and WT siblings) affected by NSC23766. 996 DEGs unique for control *tsc2^vu242/vu242^* were rescued after NSC23766 treatment (Fig. 5 E). Furthermore, the expression of almost all significantly altered gene subsets in *tsc2^vu242/vu242^*associated with actin cytoskeleton dynamics, axon growth and mTOR were normalized after treatment of Rac1 inhibitor exhibiting a marked downregulation post-treatment (Fig. 6). Together, these analyses indicate that Tsc2 loss triggers transcriptional changes converging on mTorC1 hyperactivation, Rac1-driven actin cytoskeleton remodeling, and axonal pathway dysregulation, which can be modulated by Rac1 inhibition.

**Figure 5.**
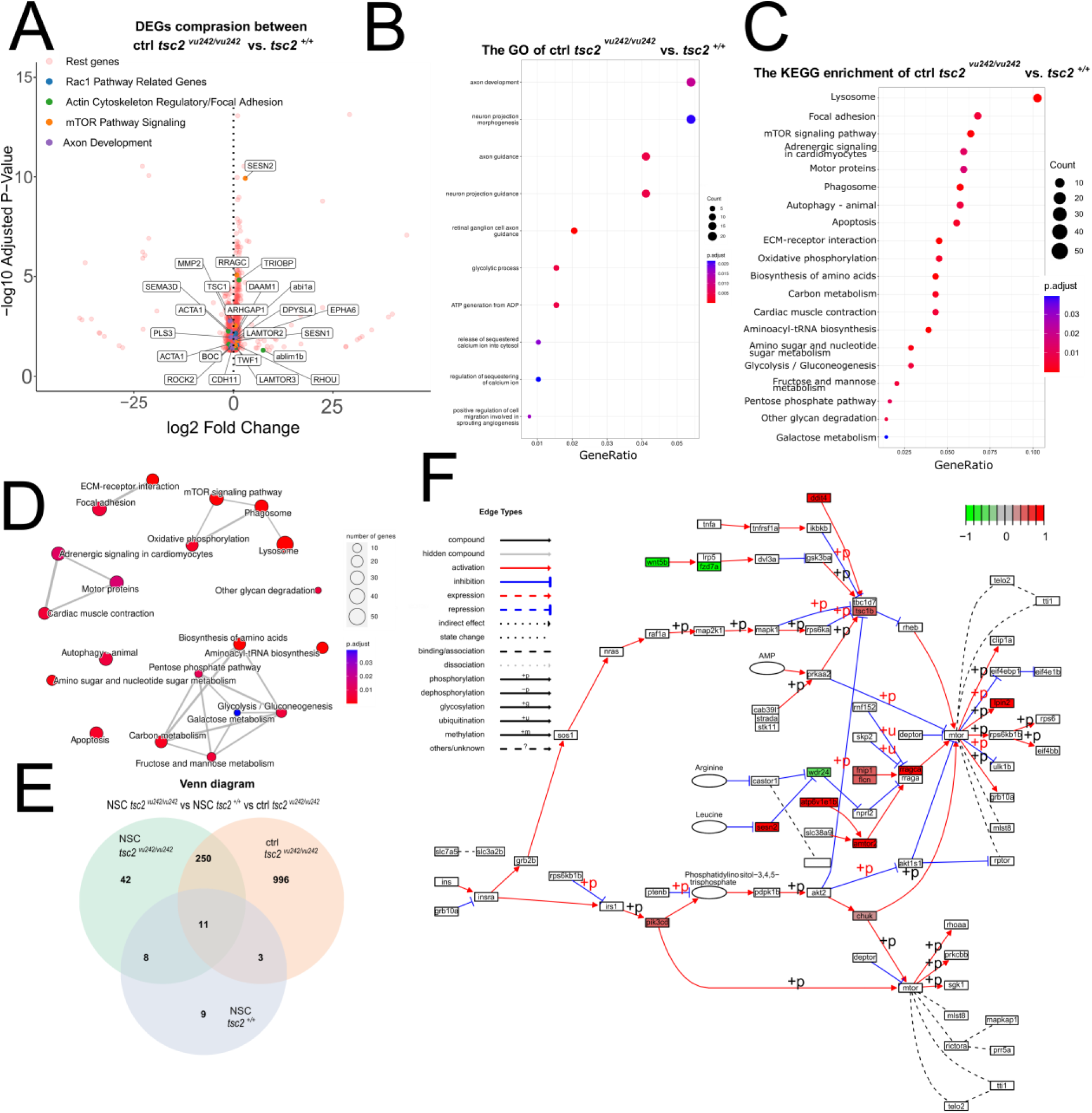
Molecular network interactions converging on Rac1 signaling. **(A)** Volcano plot of RNA-seq human orthologous data showing DEGs in Tsc2-deficient vs. wild-type brains. **(B)** Filtered GO term enrichment analysis of all upregulated DEGs between *tsc2^vu242/vu242^* and *tsc2^+/+^*zebrafish. **(C)** The KEGG pathway analysis all upregulated DEGs between control *tsc2^vu242/vu242^* and *tsc2^+/+^*zebrafish brains. (**D**) Network diagram showing shared interactors and signaling convergence based on KEGG enrichment pathway analysis. (**E**) Venn diagram comparing overlapping DEGs across Rac1 inhibition and Tsc2-deficiency. For Venn diagram 1260 DEGs founded between *tsc2^vu242/vu242^* and *tsc2^+/+^*zebrafish was used to check the significance after NSC23766 treatment in *tsc2^vu242/vu242^* vs *tsc2^+/+^* vs untreated *tsc2^vu242/vu242^*. All these variants were compared to *tsc2^+/+^.* (**F**) Detailed mTOR pathway signaling in control *tsc2^vu242/vu242^* zebrafish brain shows significant changes in fold changes of genes marked in red are significantly upregulated, whereas genes marked in green are downregulated.

**Figure 6.**
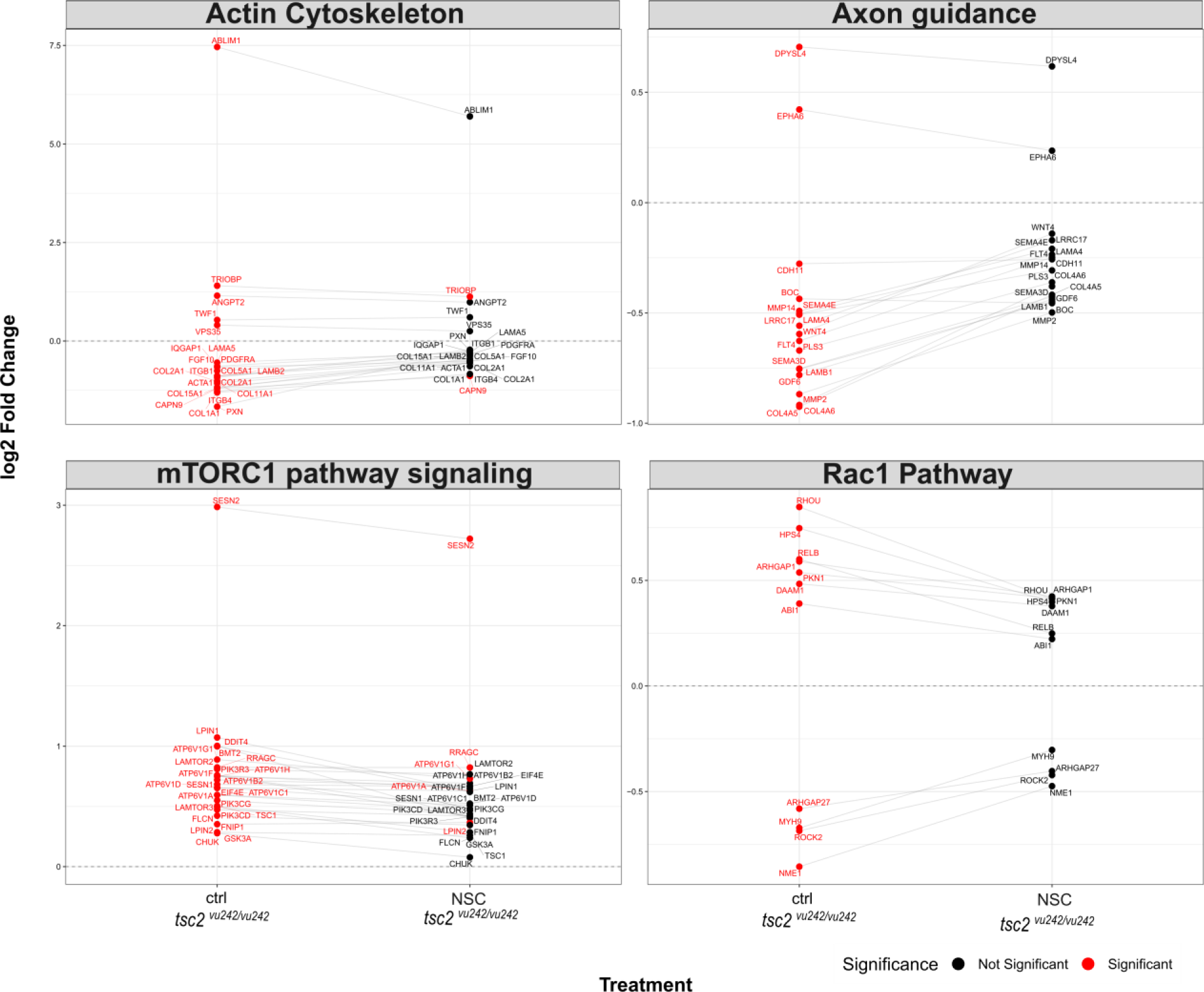
NSC23766 treatment normalizes expression of genes related to axon development, actin cytoskeleton regulation, Rac1 pathway and mTOR signaling in *tsc2^vu242/vu242^* zebrafish.

## 4. Discussion

TSC symptoms arise during brain development through mechanisms that are not fully understood. While the disease is clinically defined by the presence of benign tumors throughout multiple organs, the neurological burden extends far beyond epilepsy. TANDs are highly prevalent, often overlapping, and typically emerge during early development (Curatolo et al., 2015; de Vreis et al., 2015). Over 90% of TSC patients have TANDs which include behavioral, psychiatric, neuropsychological, and social and emotional processing issues (Krueger et al., 2013; de Vries et al., 2018). An estimated 84% of patients have seizures, although ascertainment bias likely means that a smaller percentage of TSC patients have seizures (Kingswood et al., 2014; Nabbout et al., 2019). As for anxiety disorder it is present in 9.7% of TSC patients (de Vreis et al., 2018).

Increasing evidence indicates that these neuropsychiatric traits are only partially explained by seizure activity or cortical tuber burden (de Vries et al., 2015). Neuroimaging studies show that TSC patients exhibit altered white matter microstructure, including lower fractional anisotropy, which correlates with cognitive and behavioral outcomes such as Autism Spectrum Disorder (ASD) (Peters et al., 2012). As for the anxiety, we previously pointed out that our model *tsc2^vu242/vu242^*also has alternations in white matter compartment and the model exhibits thinner anterior commissure and presents problems with axons crossing the brain midline (Kedra et al., 2020).

Clinical and preclinical studies have highlighted that TSC belongs to a broader group of “mTORopathies” — disorders unified by mTOR pathway hyperactivity and neuropsychiatric symptoms (Fang et al., 2023). In various cell types, mTOR shows its critical roles in multiple intracellular functions including mitochondrial metabolism, cytoskeleton organization, protein synthesis, and lipid metabolism (Caron et al., 2015; Laplante et al., 2009). These functions may be important for actin cytoskeleton remodeling, cell polarity, and kinase activation, specifically in neuronal cells, directly linking mTOR function to axonal growth, dendritic morphology, and synaptic plasticity. It has been shown that mTOR hyperactivation in neural precursor populations also increases the abnormalities in neuronal differentiation in mammalians. Hyperactivation of mTORC1 through the ectopic expression of constitutively active Rheb, an upstream positive regulator of mTORC1, in subventricular neural progenitor cells causes severe problems in neuronal cell migration and brain regional distribution of neuronal subtypes resulting in olfactory bulb heterotopia and circuit abnormalities (Lafourcade et al., 2013). Moreover, mTOR signaling pathway is implicated to the process of neuronal differentiation from adult neuronal stem cells as well (Romine et al., 2015; Switon et al., 2017; Zhang et al., 2014). Disruption of this finely tuned balance during early neurodevelopment can impair neurite extension, axon pathfinding, and dendritic branching, ultimately affecting the establishment of functional neuronal networks. On the other hand, TSC which is associated with over-expression of mTOR, usually exhibit impairments of the brain development and neuropsychiatric phenotypes (ASD, anxiety, intellectual disability, etc.) that do not necessarily correlate with mutation burden or cannot necessarily be explained by levels of mTOR activation. Therefore, other molecular pathways must interact with mTOR pathway to produce these phenotypes. While the central role of mTOR hyperactivation in TSC pathophysiology has been established, our results highlights the critical contribution of Rac1, a member of the Rho family of small GTPases, as a key mediator linking mTOR dysregulation to the characteristic brain connectivity impairments and neuropsychiatric disorders observed in this condition. Rac1 may perform this regulation during connectivity development as it links plasma membrane receptors with actin dynamics, dendritic spine morphology, and synaptic plasticity position. In line with our results, *in vitro* studies on Rac1-knock-out mouse showed that Rac1 regulates actin cytoskeleton in neuronal development through the Wasp family verprolin-homologous (Wave) complex (Tahirovic et al., 2010). Furthermore, Rac1 hyperactivity has been implicated in the various forms of intellectual disability (Reijnders et al., 2017; Zamboni et al., 2018), synaptic density and axon development alternation, callosal axon length reduction, and neuronal hyperexitability (Zamboni et al., 2018). Constitutively active forms of overexpressed Rac1 caused problems with commissural development (Reijnders et al., 2017). Moreover, hyperactive Rac1 guanine-exchanging factors (GEFs) Docks have been linked to various neuropsychiatric and neurodegenerative disorders, including ASD (Stankiewicz et al., 2014; Laurin & Cote, 2014). Our data demonstrate that Rac1 dysfunction in TSC extends beyond downstream effects of mTOR hyperactivation, representing a convergence point where multiple pathological processes intersect to disrupt brain development. Loss of TSC1 or TSC2 leads to constitutive mTOR activation, triggering developmental abnormalities that compromise neural circuit formation and function. Our findings suggest that this phenomenon may be mediated by aberrant Rac1 hyperactivation. Previous studies indicate crosstalk between mTOR and Rac1 pathways: TSC1 inhibits Rac1, whereas TSC2 blocks this inhibition, enabling Rac1 activation and subsequent Rho inhibition, promoting stress fiber disassembly and focal adhesion remodeling (Goncharova et al., 2004). Our study shows that Rac1 inhibition by NSC23766 and EHT1864 reduces mTORC1 hyperactivity, as indicated by decreased P-Rps6 fluorescence intensity, without significantly affecting the number of P-Rps6-positive cells (Fig. 3 A–B), suggesting Rac1 modulates mTORC1 signaling magnitude rather than activation extent.

These findings align with studies using inducible Rac1 deletion, which inhibits basal and growth-factor activation of both mTORC1 and mTORC2 (Saci et al., 2011), indicating that Rac1 directly interacts with mTOR and mediates its localization at specific membranes. Loss-of-function *tsc2* mutations increase Rac1 activity, promoting cell migration, lamellipodia formation, and reactive oxygen species production. However this was showed in renal carcinoma cells. However, this was shown in renal carcinoma cells. As for brain cells, TSC2 loss-of-function mutations may similarly increase Rac1 activity, potentially altering neuronal migration, promoting abnormal dendritic or axonal growth (Suzuki et al., 2008). TSC2 is crucial for activating RAC1 which was shown by TSC2 knockout or knockdown in fibroblast and cancer cells, and reintroduction of TSC2 or short-term rapamycin treatment rescue the defects in cell migration and and RAC1 GTPase activity (Larson et al., 2010). In Tsc1-deficient cells, expressed TSC1 suppresses proliferation and migration by reducing Rac1 activity and lamellipodia/filopodia formation, and promotes apical actin fiber formation via an mTOR-independent Rho-ROCK pathway, confirming its role in cytoskeletal organization and apical-basal cell polarity (Ohsawa et al., 2013). TSC2-deficient cells also exhibit impaired spreading, polarized cytoskeleton formation, and reduced motility.

We show that active Rac1 (GTP-bound form) was hyperactive in our *tsc2*-loss zebrafish model brains compared to WT and heterozygous siblings, which was proved by using G-LISA and FRET techniques. This confirms Rac1 dysregulation in TSC, which occurs through multiple interconnected pathways that converge to disrupt normal neuronal development and synaptic function. The loss of TSC2 protein leads to sustained mTORC1 activation, which in turn may affect Rac1 activity through several mechanisms. Abnormalities in dendritic spines and altered synaptic structure are hallmarks of TANDs which were observed in TSC animal models (reviewed in Shimada et al., 2022). Such synaptic disruptions are thought to extend beyond the level of individual neurons, ultimately affecting axon development and, consequently, the integrity of white matter compartments.

Longitudinal MRI-DTI data of TSC infants patients with ASD diagnosed at 24 months showed reduced FA across several white matter tracts compared to TSC subjects without ASD (Prohl et al., 2019). This suggests that TANDs are more likely connected with brain connectivity changes than caused by tumors and epilepsy. In line with this, our study focused on the AC, a key interhemispheric white matter tract in zebrafish. By measuring the width of the AC, we were able to assess how early disruptions in neuronal and synaptic development may translate into macroscopic alterations in brain connectivity. We already have shown before that in our *tsc2*-deficient zebrafish AC was statistically thinner compared to WT and heterozygous siblings (Kedra et al., 2020). After treatment with Rac1 inhibitors, we could observe a partial rescue of AC thickness, which suggests that abnormal Rac1 signaling contributes to structural forebrain abnormalities in *tsc2^vu242/vu242^* fish and may be modifiable through pathway inhibition (Fig. 4). This results correlate with anxiety-like behavior present in our zebrafish model which was rescued after Rac1 inhibitors treatment.

In most of the experiments conducted in this work, NSC23766 exerted a stronger influence on the Rac1 signaling pathway than EHT1864. This predominance is closely tied to their distinct mechanisms of Rac1 inhibition: NSC23766 prevents activation by blocking Rac1’s interaction with its GEFs such as Tiam1 and Trio (Levay et al., 2013). NSC23766 is relatively selective for Rac1 activation via certain GEFs and it does not strongly affect closely related small GTPases like Cdc42 or RhoA. In contrast, EHT1864 works by binding directly to Rac1 proteins and displacing bound nucleotide (GTP or GDP), thereby locking Rac1 in an inert, inactive state and preventing both the association of new GEFs and effector binding (Shutes et al., 2007). EHT1864 has high affinity for Rac1 and related Rac family members.

The transcriptional analysis of DEGs between our mutant *tsc2^vu242/vu242^*and *tsc2^+/+^*zebrafish showed 1260 DEGs and after NSC23766 treatment, 996 DEGs unique for control *tsc2 ^vu242/vu242^* were rescued (Fig. 5E). Based on these gene list GO analysis showed enrichment of genes associated with axon/neuron projection development and guidance (Fig. 5B). KEGG pathway analysis showed disturbed many pathways related to mTOR activity (Lysosome, Autophagy) and also mTOR pathway signaling (Fig. 5C). What is interesting, focal adhesion pathway is on the top of the pathways where a lot of genes were disturbed (Fig. 5C). Transcriptomic analysis confirms the pathology of our mutant rooted in the convergence of three major cellular processes: mTORC1 signaling, Rac1 signaling, and the Actin Cytoskeleton/Axon Guidance (Fig. 6). The marked dysregulation for orthologous human genes within these pathways provides a robust molecular basis for the brain connectivity impairments and neuropsychiatric disorders observed in TSC. The RNA-seq data clearly reflects the primary genetic defect in TSC, demonstrating widespread dysregulation within the mTORC1 pathway signaling genes. The overexpression of *TSC1* itself, along with upstream, modulatory sestrins (*SESN1* and *SESN2)* affects on AMP-activated protein kinase (AMPK) pathway, mTOR complexes, insulin-AKT and redox signaling pathways. By regulating these pathways, Sestrins are thought to help adapt to stressful environments and subsequently restore cell and tissue homeostasis (Kim et al., 2021). In our RNAseq we found many others upregulated genes that encode proteins which acts as a negative regulator of mTORC1 by activating the TSC1-TSC2 complex, like *DDIT4* (Du et al., 2018). Therefore, these findings validate the model of constitutive mTOR activity, which then sets the stage for the dramatic alterations observed in cytoskeletal and Rac1-mediated processes. The most direct molecular evidence for brain connectivity impairment comes from the massive number of dysregulated genes in the Actin Cytoskeleton and Axon Guidance pathways (Fig. 6). Disruption of these pathways aligns strongly with previously reported neural network connectivity defects in TSC patients and TSC animal models. Consistent with our findings, multiple independent studies have demonstrated aberrant neuronal wiring in TSC—arising through both mTOR-dependent and mTOR-independent mechanisms—further supporting the notion that loss of TSC1-TSC2 leads to profound cytoskeletal and axon guidance dysfunction (Catlett et al., 2021; Choi et al., 2008; Knox et al., 2007;).

In conclusion, our findings establish Rac1 as a critical molecular hub linking mTOR dysregulation to the characteristic brain structure impairments and neuropsychiatric disorders observed in TSC. At the molecular level, loss of Tsc2 function destabilizes the balance of mTORC1–Rac1 crosstalk, resulting in Rac1 hyperactivation and downstream disruption of actin cytoskeleton dynamics, axon growth and guidance, and neuronal connectivity. This dysregulation is clearly reflected in our RNA-seq datasets, which reveal coordinated disturbances across gene networks governing axon pathfinding, cytoskeletal remodeling, and Rac1/mTOR signaling, offering a multilayered mechanistic explanation for core TAND features. Importantly, this work extends beyond molecular correlations by demonstrating functional reversibility of Rac1-driven pathology. In *tsc2*-deficient zebrafish mutants, we show that Rac1 is consistently overactive, leading to impaired formation of AC as well as heightened anxiety-like behaviors. Pharmacological Rac1 inhibition not only normalizes Rac1 activity, but also rescues AC defects and rescues anxiety-like behaviors, indicating that Rac1 overactivation is not merely a correlate of TSC neuropathology but a causal and targetable driver of structural and behavioral abnormalities. Taken together, these findings highlight Rac1 as a mechanistic bridge linking genetic mutations in TSC to neurodevelopmental trajectory failures and behavioral symptoms.

## 5. Acknowledgements

We thank Kevin Ess (Vanderbilt University) for tsc2^vu242^; the International Institute of Molecular and Cell Biology (IIMCB) Zebrafish Core Facility for assistance with breeding and setting up the fish; Karim Abu Nahia (Laboratory of Zebrafish Developmental Genomics, IIMCB) for assistance and consulting in RNA-seq analysis; the IIMCB MCF for sharing the Lightsheet Z.1 (part of IN-MOL-CELL Infrastructure, which was funded by the European Union – NextGenerationEU under National Recovery and Resilience Plan); Michiyuki Matsuda for pEYFP-Rac1-ECFP plasmid (known as Raichu-Rac1/Rac1-CT); Jacek Jaworski (IIMCB) for Torcar Ypet I mCerulean plasmid; Genomics Core Facility CeNT UW (Europe, Poland) for RNA sequencing. This work was supported by the OPUS Grant 2020/37/B/NZ3/02345 from National Science Centre, Poland (to JZ).

